# PiRATeMC: A highly flexible, scalable, and affordable system for obtaining high quality video recordings for behavioral neuroscience

**DOI:** 10.1101/2021.07.23.453577

**Authors:** Samuel W. Centanni, Alexander C.W. Smith

## Abstract

With the recent development and rapidly accelerating adoption of machine-learning based rodent behavioral tracking tools such as DeepLabCut, there is an unmet need for a method of acquiring video data that is scalable, flexible, and affordable. Many experimenters use webcams, GoPros, or other commercially available cameras that are not only relatively expensive, but offer very little flexibility over recording parameters. These cameras are not ideal for recording many types of behavioral experiments, and can lead to suboptimal video quality. Furthermore when using relatively affordable commercially available products, it is a challenge, if not impossible, to synchronize multiple cameras with each other, or to interface with third-party equipment (for example, receiving a simple trigger to simultaneously start recording, or acting as a microcontroller for closed-loop experiments). We have developed an affordable ecosystem of behavioral recording equipment, PiRATeMC (<u>Pi</u>-based <u>R</u>emote <u>A</u>cquisition <u>Te</u>chnology for <u>M</u>otion <u>C</u>apture), that relies on Raspberry Pi Camera Boards that are able to acquire high quality recordings in bright light, low light, or dark conditions under infrared light. PiRATeMC offers users control over nearly every recording parameter, and can be fine-tuned to produce optimal video data in any behavioral arena. This setup can easily be scaled up and synchronously controlled in clusters via a self-contained network to record a large number of simultaneous behavioral sessions without burdening institutional network infrastructure. Furthermore, the Raspberry Pi is an excellent platform for novice and inexperienced programmers interested in using an open-source recording system, with a large online community that is very active in developing novel open-source tools. It easily interfaces with Arduinos and other microcontrollers, allowing simple synchronization and interfacing of video recording with nearly any behavioral equipment using GPIO pins to send or receive 3.3V or 5V (TTL) signals, I2C, or serial communication.

## INTRODUCTION

Preclinical studies have long relied on traditional predetermined activity patterns to assess behaviors such as affect, motivation, cognitive function, memory, motor coordination, etc. While these historical approaches have led to countless discoveries, the evolution of behavioral paradigms has yielded increasingly complex interpretations of the outcomes. Biased behavioral scoring (i.e. the need for *a priori* knowledge of behaviors to score) overlooks potentially unique behaviors that may occur in specific groups or subgroups within experiments. Moreover, manual behavioral scoring is highly vulnerable to human error, biases, and low inter-rater reliability. Using unbiased pose estimation techniques with open-source machine learning-based software such as DeepLabCut^1, 2^, and behavioral mapping or clustering analysis with packages such as B-SOID^3^ or VAME^4^ have begun to shift behavioral neuroscience into a new era of behavioral analysis and categorization. These techniques are able to segment behaviors in an unbiased way, and eliminate inconsistencies due to human error and inter-rater variability. To maximize the potential for these and other programs, experimenters must be able to capture high resolution videos. Moreover, the ability to interface recording equipment with real-time controllers for third-party data collection equipment, such as *in vivo* measurement of brain activity (e.g. fiber photometry/miniscopes), optogenetic LED drivers, or MedAssociates® chambers unlocks the potential for closed-loop experiments, and very easy time-locking of video recordings with data from these systems.

Here we introduce PiRATeMC (<u>Pi</u>-based <u>R</u>emote <u>A</u>cquisition <u>Te</u>chnology for <u>M</u>otion <u>C</u>apture), an affordable, user-friendly, modular, open-source camera system that runs off a Raspberry Pi (RPi) single-board computer and an accompanying 8-megapixel (MP) Camera Board (Sony IMX219 CMOS sensor). These cameras can record high quality video under either infrared (IR) or white light. Moreover, they offer far more flexibility in recording parameters than most commercially available camera systems. The user can manually set recording parameters like frame size, frame rate (up to 120FPS), white-balance (crucial for high-quality IR videos), brightness/contrast, ISO, saturation, bitrate, and many more (details in **Table 2** and **Supplemental Table 2**). Finally, the PiRATeMC system can easily be controlled remotely via *ssh* (remote secure shell) over a local area network (LAN), and clusters of cameras can be controlled synchronously with millisecond precision using ClusterSSH (easily installed via the advanced packaging tools (apt) in Linux or Homebrew in MacOS), allowing either multi-angle recording of single subjects (e.g. for 3D-DeepLabCut^5^), or an easy method of recording a large number of behavioral sessions simultaneously. We provide step-by-step instructions to physically assemble the camera and Raspberry Pi, as well as a cloned operating system (PiCamOS) that can be uploaded to an SSD card (source code for building PiCamOS is also available). Lastly, we describe a simple data management pipeline, whereby with minimal user modification of PiCamOS, a large number of RPiCams can be controlled simultaneously, and deposit all recorded videos in the same ‘sink’ directory on a remote local network (LAN) host. The goal of this paper is to provide an easily obtainable recording system that is highly flexible in recording parameters, can interface with numerous real-time equipment controllers and other open-source analysis software, and can easily be scaled up to record a large number of videos simultaneously. Increasing accessibility and usability of acquisition and analysis programs for assessing rodent behavior has the potential to provide a wealth of new behavioral data that may have been overlooked using traditional approaches.

**Table 1.** Parts List. This table contains the minimum essential parts to get started with one camera. Only one camera (either NoIR or standard camera board) is needed. For mixed dark/white light use, we recommend the NoIR camera. For a full list of optional parts/accessories, as well as notes about each component, see **Supplemental Table 1**.

| **Basic Camera Necessities:** |  |  |  |
| --- | --- | --- | --- |
| **Part:** | **Vendor (suggested):** | **Product #:** | **Price:** |
| Raspberry Pi 3 Model B+ (or newer) | Adafruit Industries, Newark Electronics | 3775 (Adafruit), 49AC7637 (Newark), | \$35.00 |
| Raspberry Pi NoIR Camera Board v2 - 8MP | Adafruit Industries | 3100 | \$29.95 |
| Raspberry Pi Camera Board v2 - 8MP | Adafruit Industries | 3099 |  |
| 5V 2.5A Switching Power Supply | Adafruit Industries | 1995 | \$7.50 |
| Kingston 32GB microSD card | CDW | 5849358 | \$4.99 |
| Flex Cable for Camera (various lengths available) | Adafruit Industries | 1730 | \$2.50 |
| **Total:** | | | **\$79.94** |

**Table 2.** Abbreviated list of Raspivid commands that can be used in recordVideo.sh line 3.

| Short form | Long form | Explanation |
| --- | --- | --- |
| -? | --help | Help |
| -w | --width | Width of image (default 1920) |
| -h | --height | Height of image (default 1080) |
| -o | --output | Output filename |
| -fps | --framerate | Number of frames to record per second |
| -sh | --sharpness | Image sharpness (-100 to 100) |
| -co | --contrast | Image contrast (-100 to 100) |
| -br | --brightness | Image brightness (0 to 100) |
| -sa | --saturation | Image saturation (-100 to 100) |
| -awb | --awb | Set auto white balance (options= off, auto, sun, cloud, shade, tungsten, fluorescent, incandescent, flash, horizon, greyworld) |

## METHODS

Note that any text highlighted in grey below represents inputs or outputs from the Linux terminal, and is case-sensitive.

### Camera Assembly

Raspberry Pis (RPis) are single-board computers that run a full Linux operating system (Raspbian), and thus offer greater computational capabilities than other single-board computing devices such as Arduinos. For a simple recording setup, we recommend using the RPi 3 Model B+, as this model is the last to have a standard HDMI output port and micro-USB charger. RPi 4 and newer can certainly be used, however newer models only have micro-HDMI output, and a USB-C power input, which likely necessitates purchase of additional adapters. There are two options for 8MP camera boards that connect to the RPi camera interface: a NoIR Camera Board, and a standard Camera Board v2. The naming of these cameras is not intuitive, the NoIR camera is the camera that <u>is</u> sensitive to infrared light, (i.e., does <u>not</u> have an IR filter), but may produce lower quality videos under white light. If you only plan to record experiments under white light, the standard Camera Board v2 is recommended, as an IR filter will enhance video quality under white light. In order to set up a single camera, the minimal parts required are listed in **Table 1**. Note that only one of the two camera options (NoIR or standard) is necessary, and these cameras come with a Flex Cable, however the stock cable is only 6” long, so while a longer Flex Cable is not absolutely necessary, we consider a longer flex cable necessary equipment, and the price is very low.

Once the required parts have been gathered, assembly is very easy. First, replace the stock 6” camera cable with a longer one with a length of your choice (we use 24” for recording inside operant chambers). When inserting flex cables, the silver leads of the cable always face away from the slider clamp that secures the cable in place, on both the camera and the RPi. Instructions and example photos for attaching the camera to the RPi shown in **Figure 1**.

**Figure 1.**
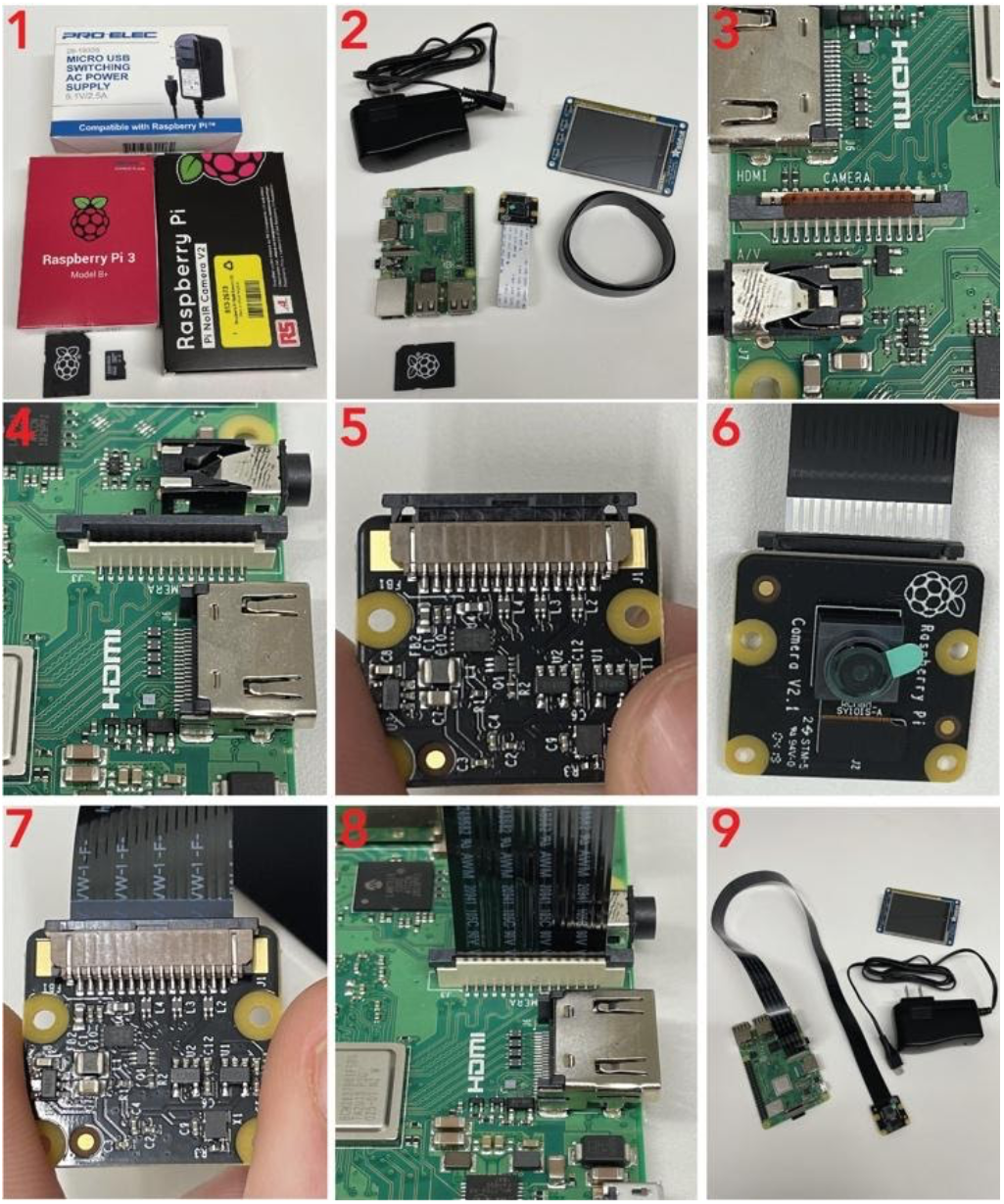
Attached the Camera to the Raspberry Pi. Panel 1 shows the minimum parts needed for assembly. Panel 2 shows the unpackaged parts, as well as the optional 24” camera cable, and PiTFT touch screen (see Supp. Table 1 for details). The camera cable is attached at both ends by black clamps. Remove the plastic covering from the Pi (Panel 3), and gently lift the edges as shown in Panel 4. If you are replacing the stock camera cable with a longer cable (recommended), optionally do the same to the clamp on the camera, and remove the cable as shown in Panel 5. To attach a new camera cable, the silver leads on the cable face the same direction as the lens (Panel 6). Insert the cable all the way into the camera, then gently press the clamps back down (Panel 7). Follow the same procedure to insert the new cable into the camera port on the Pi, with the silver leads facing away from the clamp (towards the HDMI port on Pi 3), then gently press down on the edges to secure the cable in place. Panel 9 shows an assembled Pi with a 24” camera cable.

### Remote Controller Hardware Configuration

While RPis can be connected to a keyboard and monitor and operated as standalone computers, the most convenient way to control the cameras is via a remote secure shell (SSH) connection. This allows the RPi to be controlled from a remote computer (referred to here as the ‘remote controller’) connected to the same network as the RPi, and the remote controller can also serve as a data sink where videos are automatically transferred after recording. Because the RPi operating system is a Linux distribution (Raspbian), it is most convenient if the remote controller is also running a UNIX-based operating system (Mac OSX, Ubuntu, or any Linux Distribution). Users that only have Windows PCs available should use a Bash emulator such as Git Bash (https://gitforwindows.org/), which will allow use of all terminal commands described for UNIX based operating systems below (see Supplemental Material - Notes for Windows Users for details). If you plan to scale up the number of RPis and cameras for larger experiments, we recommend using a mini-PC such as the AWOW MiniPC running Ubuntu 18.04 or later (see **Supplemental Table 1** for product details). Because many institutional IT departments prohibit use of network switches, the remote controller should have at least a dual-port network adapter card, as transmitting video data over WiFi is orders of magnitude slower. This will allow the user to maintain connectivity to the internet and backbone institutional LAN on the remote controller using one network port, and also to configure the remote controller to act as a DHCP server, which creates its own LAN to manage the RPis. Detailed instructions for this process are at https://github.com/alexcwsmith/PiRATeMC/tree/master/docs/networking, and a schematic of this architecture is shown in **Figure 2**. A network switch can then be connected to the second port, allowing the remote controller to assign and manage IP addresses to any RPis connected to the switch, without the RPis having access to the main institutional network. Thus, PiRATeMC does not pose any security risk at hospitals/university medical centers and other institutions where security is a significant concern. Detailed instructions for configuring this network are provided in supplemental materials, and an instructional walk-through video can be found in the documentation in the GitHub repo linked above. The packages that need to be installed (via sudo apt install) on the remote controller are: *openssh-server, clusterssh, isc-dhcp-server, bind9*. Before beginning any reconfiguration, users should connect the remote controller to the institutional backbone network, and enter ip a s into a terminal, and record the information there (or write to a plain text file with echo $(ip a s) > ∼/Desktop/ipas.txt), and also create backup copies of the files at: /etc/netplan/*.yaml and /etc/dhcp/dhcpclient.conf, and record the institutional network’s default network settings such as domain name servers (cat /run/resolve/resolv.conf > ∼/Desktop/DefaultDNS.conf) and gateway (echo $(ip -a route) > ∼/Desktop/ipRoute.conf). Users planning to record videos with only one camera should not have an issue connecting to the institutional network without an intermediate network switch, or in the simplest use case, can connect a PiTFT Touch Screen (add on accessory, see **Supplemental Table 1**) or monitor and a keyboard directly to the RPi and record videos locally without any network connectivity.

**Figure 2.**
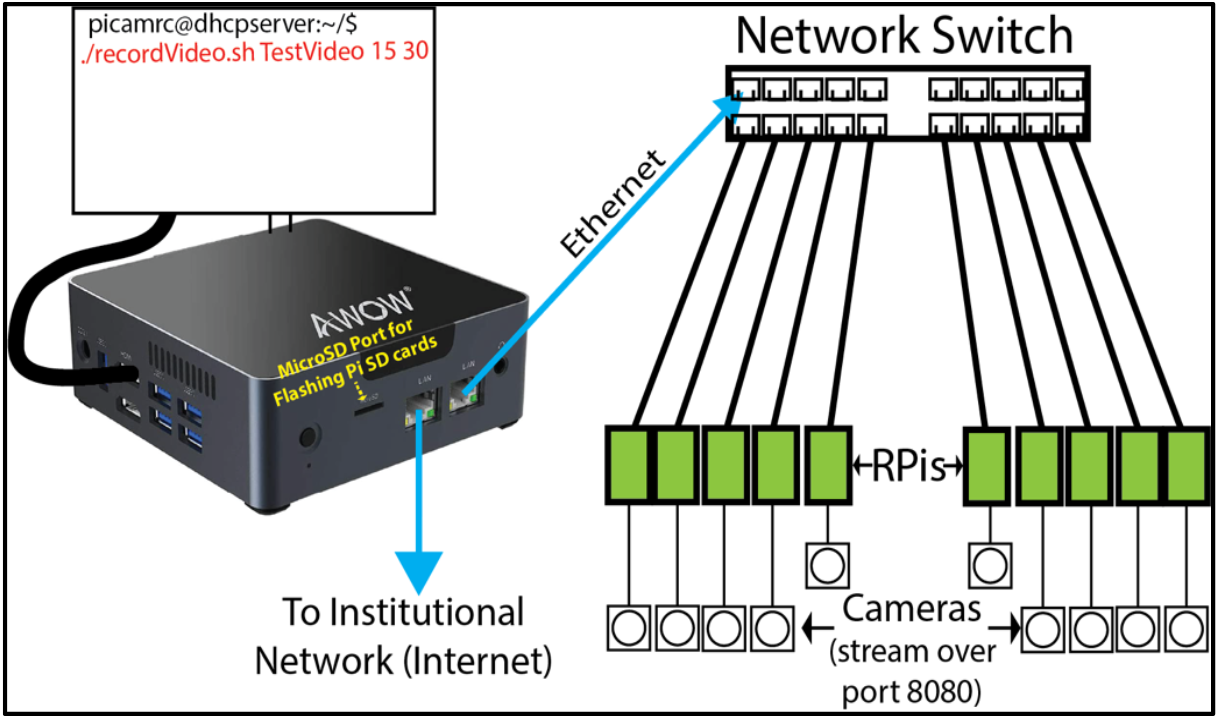
Schematic showing infrastructure for creating isolated LAN separate from main institutional network to allow using an affordable unmanaged network switch to manage clusters of PiRATeMC Cameras. Many institutions (every single one we personally have dealt with, though these have all been hospitals/medical centers) prohibit these network switches on their networks, though you may try plugging a network switch directly into your institutional network, and if it is allowed to connect you do not need to configure your own isolated LAN.

### Software Configuration

Two versions of pre-configured and fully functional cloned disk images of the PiRATeMC operating system are available to download from Google Drive (along with other resources) at: https://tinyurl.com/jaaeu8s2. The ‘standalone’ version is optimally suited for users who want to use a single camera and are able to directly connect the RPi to their institutional LAN, or who will use an RPi without a remote connection. The ‘cluster’ version is optimized for users who want to build clusters of cameras to record a large number of behavioral sessions simultaneously using a network switch and an isolated LAN. The source code for these operating systems, and documentation is available on GitHub at https://github.com/alexcwsmith/PiRATeMC. To flash the operating system onto the SD card that will be used as the storage drive of the RPi, use rpi-imager, available for download at https://www.raspberrypi.org/software/, or by typing: sudo apt install rpi-imager into a Linux terminal. We have created a video showing detailed steps for writing PiCamOS to the SD card and for editing system variables described below. On our system this process takes ∼3 minutes, you can find this video here: https://youtu.be/Ier37fRuUos. PiCamOS is a modified version of Raspbian Buster, and comes pre-installed with drivers for the PiTFT Resistive Touch Screen, the UV4L video streaming service, and WebRTC. PiCamOS also has pre-installed shell scripts for logging IP addresses to the remote controller at startup (sendIP.sh), and for recording videos with highly flexible parameters (recordVideo.sh).

After flashing PiCamOS to the SD card, it will be automatically unmounted/ejected. Before placing the SD card into the RPi, there are 4 variables to change in a file called .bashrc that will allow the RPi to communicate with the remote controller the first time it is powered on. To edit this .bashrc rile, re-mount the SD card to the remote controller, and navigate (cd) to the folder /rootfs/home/pi/ on the SD card (on Ubuntu/Debian Linux this is likely mounted at /media/*<YourComputerName>*/rootfs/. If you don’t know the nickname of your computer, simply cd /media/ then type ls, which will likely only display one folder (the the name of your computer), and cd into that folder, where ls will show rootfs mounted). On Raspbian (and all UNIX-based operating systems), any file names that start with ‘.’ are hidden files, and can be viewed with ls -a, or if using a GUI, make sure that hidden files are viewable (Ctrl+h on Linux, Cmd+Shift+. on Mac). Open the file ‘.bashrc’ with either a GUI text editor (gedit or TextEdit), or with a terminal text editor like *nano* or *vim* by typing into the terminal: nano .bashrc. This .bashrc file is a type of configuration script (equivalent to .bash_profile on Mac OSX) that is run every time a new terminal is opened, and defines system variables, runs startup scripts, or performs other tasks to initialize a user session. Near the top of this file you will see four lines beginning with ‘export’ that define system variables to edit to be valid with your system:

1. export REMOTEPATH=rpicam@10.1.1.243
  a. This is the username & IP address to an account on your remote controller computer that also serves as a ‘data sink’. This information will be used for automatically transferring recorded videos from the source (RPi) to the sink (remote controller). If you are setting up a network for a cluster and using the provided network files, this IP address is the default IP address for the remote controller, so you only need to change the username.
2. export REMOTEPASS=$(cat .pass_file)
  a. This variable reads the password to the REMOTEPATH account from another hidden file in the /home/pi/ directory, called .pass_file. Note in a graphical interface the .pass_file command will appear locked, and is only viewable by the file owner or the system root. In order to edit the contents of this file, open it with sudo nano .pass_file in a terminal while in the /home/pi/ directory.
3. export REMOTELOGPATH=$HOME/RPi_Sessions/
  a. Path to a directory on the remote controller to write log files upon RPi startup. Make sure this folder exists on the remote controller.
4. export REMOTEVIDPATH=/d1/studies/Raspicam/
  a. Path to a directory on remote computer to store recorded video files. Again, make sure this folder exists on the remote controller.

Finally, you (optionally) can change the RPi hostname to something unique and recognizable at this point (the default is raspberrypi; we use the room and box # of the chamber each RPi is recording from), or you can change the hostname after you have established a connection over ssh. You can change the hostname in the rootfs directory before starting up the RPi for the first time with sudo nano /media/<YourComputerName>/rootfs/etc/hostname, which contains only the hostname, and sudo nano /media/<YourComputerName>/rootfs/etc/hosts, where you will find the hostname after the string 127.0.1.1, likely at the bottom of the file. **Caution:** The hostname must be identical in these two places, if it is not, you will see the warning *“sudo: temporarily unable to resolve hostname”* when the Pi is booted. If you see this warning after connecting to the Pi, check that the hostname is identical in those two files directly on the RPi via sudo nano /etc/hostname and sudo nano /etc/hosts.

After the system variables and optionally the hostname have been set, plug the RPi into an ethernet port, either on a network switch if running in a cluster, or directly if running on a main institutional backbone network. If everything is configured correctly, shortly after powering on, the RPi will transmit a log file to the directory set in $REMOTELOGPATH above. If running in a cluster, typing dhcp-lease-list on the remote controller will output a list of connected RPis and their IP addresses. You can then access the RPi from a terminal on the remote controller via ssh (the default password is raspberry). Note if the RPi does not have internet connectivity, it may record the wrong date/time. Once connected via ssh, the date/time can be set from the terminal with sudo date --set “YYYY-MM-DD HH:MM:SS”. Note that once you are able to connect to the RPi via ssh, you can also move files to/from the RPi and any computer connected to the same network with the scp command, for details input man scp into the terminal.

### Interfacing with third-party equipment

The RPi has 40 general purpose input/output (GPIO) pins that are capable of sending and receiving 3.3V and 5V (TTL) signals, inter-integrated circuits (I2C), and serial communication. These offer a simple method of interfacing with Arduinos or other microcontrollers, or for using the RPi as a microcontroller itself. There is a large online community dedicated to discussing novel methods of using both RPis and Arduinos as microcontrollers for hundreds of applications across scientific disciplines, as well as hobbyists using these devices for at-home DIY engineering projects. Thus, there are several pre-made chips and ‘HATs’ (Hardware Attached on Top) so that users do not need any experience with electrical engineering to accomplish relatively simple tasks like using the RPi to control an LED driver, a light/tone generator for fear conditioning, an ultrasonic vocalization microphone, or even a motorized peristaltic pump. Though we have not included detailed instructions for these uses in this version of the manuscript, we hope that the neuroscience community will build upon this system and continue to make novel applications open-source, and we will elaborate further on these capabilities in future versions. See the “Optional Microcontroller Interfacing Equipment” section of **Supplemental Table 1** for a small list of parts we believe may be of particular interest. **Figure 3** shows a pin map of the RPi 3B+ GPIO pins to give a better idea of the types of interfacing and communication that are possible.

**Figure 3.**
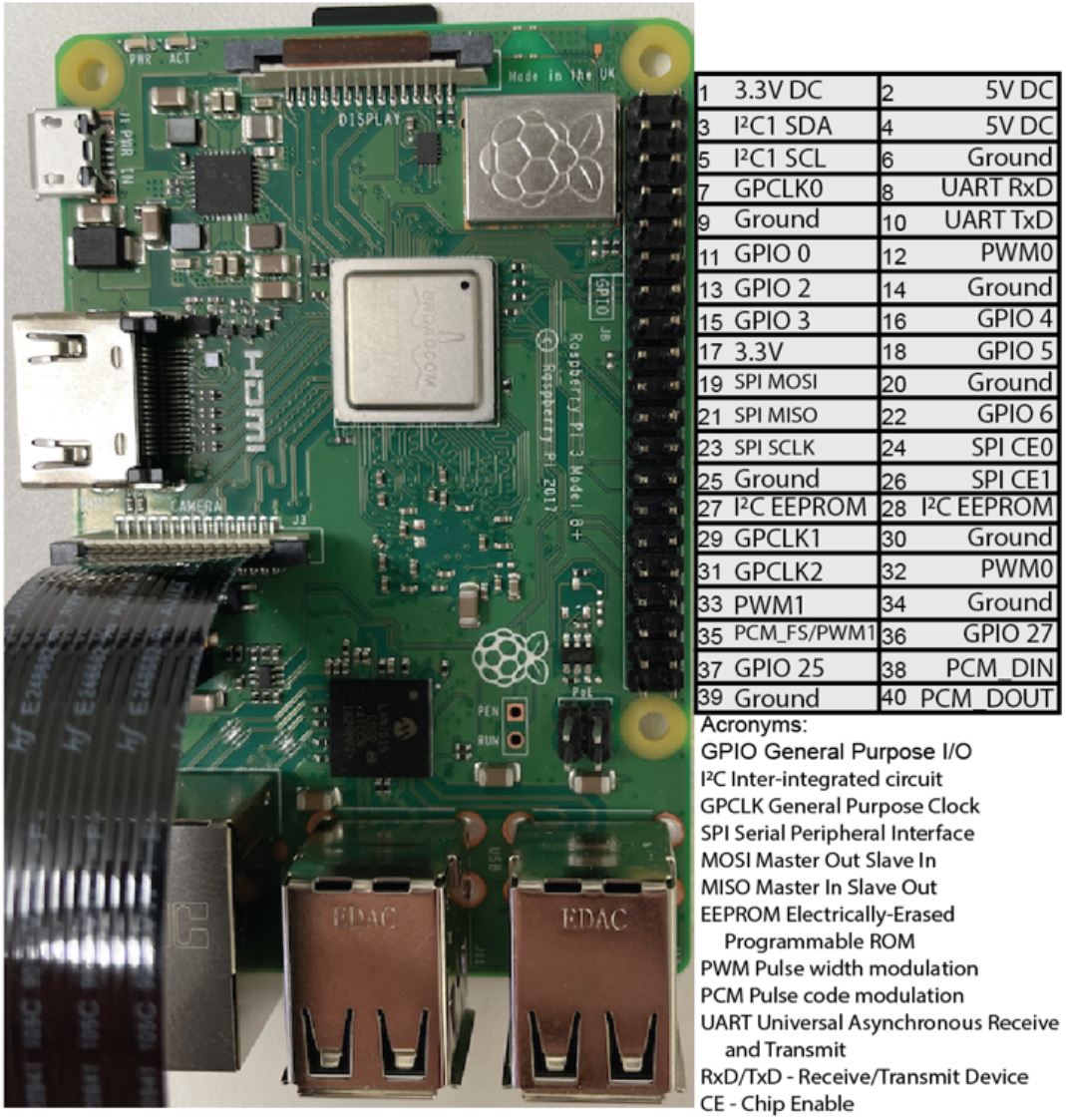
GPIO Pin Diagram for Raspberry Pi 3 Model B+.

## RESULTS

### Recording or Streaming Video

After the PiRATeMC hardware and software have been configured, and you are able to access the RPi through an SSH connection, you are nearly ready to begin recording or streaming videos. The first time you power on each RPi, the first command entered should be sudo raspi-config. This will bring you to a GUI configuration manager, where it is advised to change the password, the hostname (if you did not change it in the rootfs while flashing the SD card), and the location/time zone settings. If you are based outside of the USA, it may be necessary to change the keyboard layout as well. There are several methods for recording or streaming videos.

### Recording via recordVideo.sh script

The recordVideo.sh script is the most powerful method for recording high-quality videos, as it offers the most flexibility of recording parameters. The usage for running this script is:

- ./recordVideo.sh <VideoName> <DurationInMinutes> <FramesPerSecond>

For example, ./recordVideo.sh TestVideo1 0.25 30 will record a 15-second video with 30 frames per second, and will be saved with filename TestVideo with the hostname and date appended to the filename, in a .mp4 format. This video will be stored locally on the Pi, and will also be automatically transferred to the location set in the $REMOTEVIDPATH variable above. Please note that videos will also be saved locally on the RPi, and should therefore be deleted regularly to avoid running out of storage. **<u>This is critical</u>**, as full storage will not prevent the recording from starting, only from saving at the end of the recording. To clear all videos from the RPi, enter the command rm *mp4 while in the home folder. **Warning:** use caution when combining the rm command with the wildcard *, as an accidental space between the * and mp4 can delete the entire OS, forcing you to reinstall. A safer, but more time-consuming way is to use the rm command with the exact filename (e.g., rm TestVideo1.mp4), or to use a shared string within video names (e.g. rm Test*.mp4). To confirm the files are removed, use the command ls *.mp4. No files with an .mp4 extension should be present.

**Table 2** shows some of the most useful recording parameters that can be manually set. To change these parameters, edit the recordVideo.sh script with nano recordVideo.sh, and parameters can be inserted in line 3 after the raspivid command. The impact of some of these recording parameters on video quality is shown in **Figure 4**. A caveat of the minimal setup we have described here is that the recordVideo.sh script does not easily allow simultaneous viewing of the video while it is being recorded. This can be remedied by attaching a 2.8” touchscreen to the RPi and running the command: sudo python3 ∼/setupScripts/adafruit-pitft.py, and following the prompts. The final prompt of the setup process will ask “Would you like the console to appear on the screen?”, and answering “No” to this will result on the video being displayed on the screen while it is recorded. Alternatively, a 7” display can be purchased and attached via a flex cable to the Display Port of the RPi, and this will display the stream while recording is ongoing. Part numbers for both options are provided in **Supplemental Table 1**. As a third option, the RPi can be plugged directly into a monitor via the HDMI port, and the monitor will display video while it is being recorded. We do not consider these options necessary, and they increase the price of each PiRATeMC unit by 2-3 fold, and in our experience while recording many videos simultaneously inside of operant boxes, there is no real need to be view the streams live, as videos are analyzed with DeepLabCut immediately following acquisition (potentially automatically in future versions, see discussion). A fourth option to view videos while recording is via online streaming with WebRTC (described in detail below). While this adds no additional cost, WebRTC offers far fewer parameters to change, and can cause inconsistent recordings (i.e., dropped frames).

**Figure 4.**
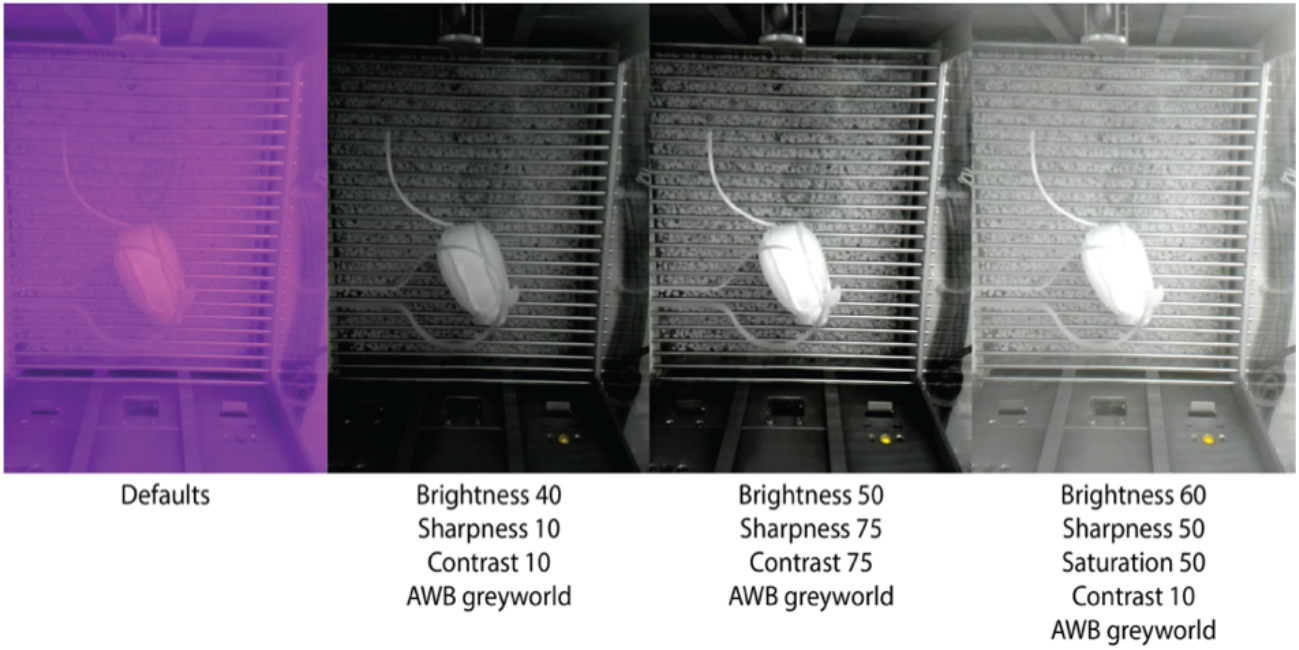
Demonstration of the effect of different recording parameters on video quality under infrared light. Most obviously, setting the auto-white-balance (--awb option) to ‘greyworld’ is necessary for correcting for the red-shifted light from the infrared light source. Tuning brightness, saturation, sharpness, and contrast also have obvious effects on video quality. See Table 2 for the parameters we find most helpful, and Supplemental Table 2 for all available options.

### Streaming via UV4L

You can also stream videos over the network from the remote controller (or any computer connected to the same network as the RPi) through port 8080 by opening your internet browser of choice, and entering the RPi IP address followed by :8080 into the address bar, e.g. 10.1.1.50:8080. This will take you to a UV4L Streaming Service configuration page shown in **Figure 5**.

**Figure 5.**
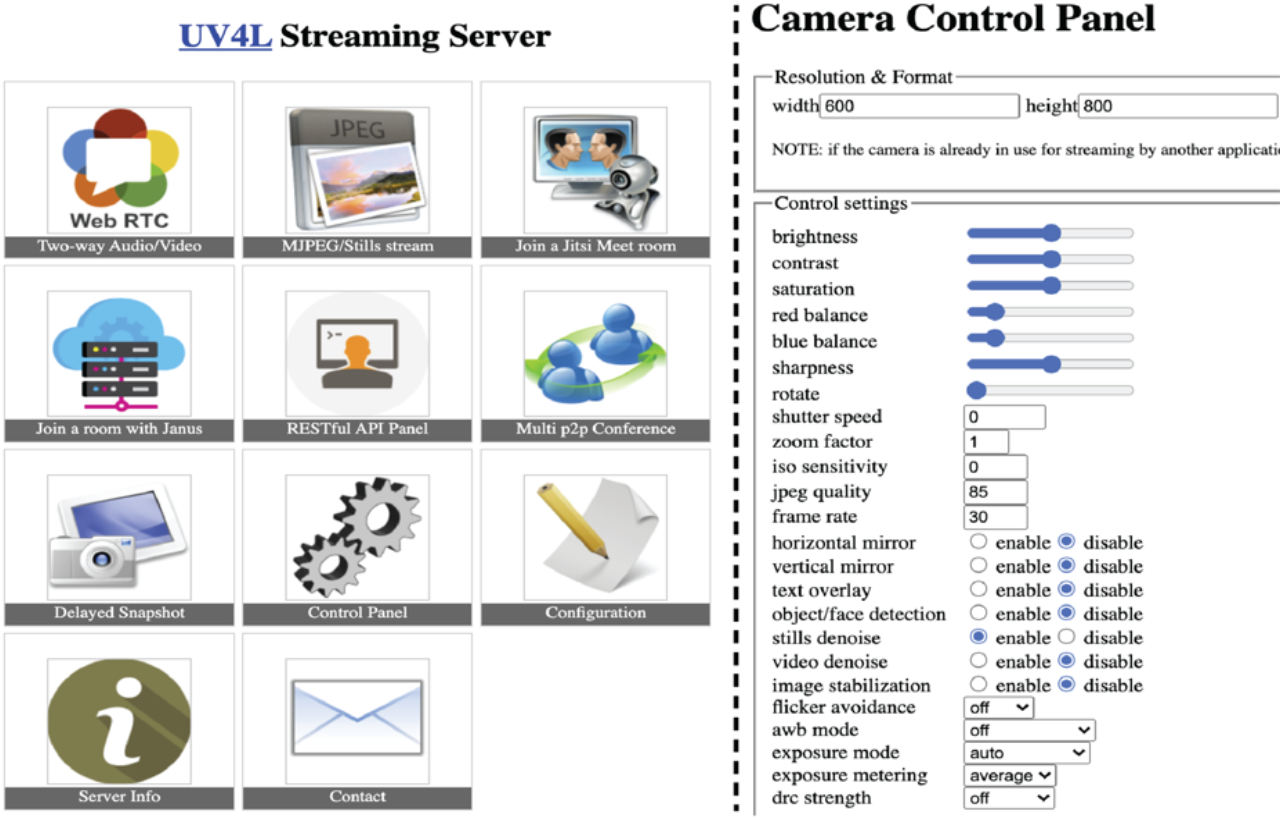
UV4L Streaming Server Control Pages. The left side is the UV4L home page. On the right is the destination of the ‘Control Panel’ link. Clicking ‘MJPEG/Stills stream’ on the UV4L home page results in a live stream (not shown).

The UV4L streaming service is ideal for optimizing recording parameters to use in the recordVideo.sh script, and for viewing the live stream while fixing cameras in place to optimally capture the field of view for behavioral equipment. However, the UV4L service does not offer a way to both view the live stream and record simultaneously, and it does not offer a way to synchronize multiple cameras. The UV4L driver also does not offer a ‘greyworld’ mode for the AWB setting, so video quality in these streams will not be as good as can be accomplished via the recordVideo.sh script. However, this does offer a powerful way to optimize camera position, and to test the effect of manipulating parameters such as brightness/contrast/saturation on video quality.

### Streaming via WebRTC

Similar to streaming via UV4L, you can also stream through an internet browser by entering an IP address followed by :8080/stream/webRTC/. This method does allow simultaneous recording and streaming, however it does not offer flexibility or tuning of recording parameters (though there are several presets to choose from for frame size/rate), and is more prone to dropped frames. Using this method, videos will be recorded and saved locally on whatever computer is running the web browser, not necessarily to the path saved as the data sink.

## DISCUSSION

A major confound in behavioral neuroscience is the rigidity and inflexibility of many commercially available behavioral tracking software, in addition to the high costs of implementing these systems. Many programs require a specific type of camera, offer limited options for adjusting the image, and lack the adaptability to sync across other acquisition programs. Moreover, specific video file outputs require time consuming conversion software to analyze videos with external programs. Here we present an affordable, scalable, customizable, open source video recording configuration, PiRATeMC, that can be optimized for recording animal behavior in any setting. Setting up PiRATeMC requires very little programming/coding knowledge. Simply upload the software onto a microSD card, set four environment variables on the microSD card while inserted into a PC, insert it into the RPi and power on, and record videos. Video recording settings can then be modified to get very high quality videos of large behavioral arenas illuminated by either infrared light or white light. Unbiased behavioral mapping is becoming increasingly important and informative in neuroscience, and the output videos from PiRATeMC can be easily incorporated into widely used open-source analysis pipelines such as DeepLabCut, B-SOID, VAME, etc.

Setting up PiRATeMC requires very little programming and coding knowledge, though experience with the Linux operating system and using the command line interface is a plus. Simply upload the software onto a microSD card, set system variables to detect the remote controller, turn on the Raspberry Pi, adjust settings, and record videos. All of the steps can be completed using simple commands in the terminal, and we attempt to clearly state which parts of the pipeline require specifying the user’s computer/network information. To provide support beyond this paper, we include links to a GitHub, YouTube tutorials, and webpages relevant to PiRATeMC. Accordingly, we present a ground level version of PiRATeMC, and encourage those with experience in Linux coding and Raspberry Pi computing to build more code into this pipeline as needed such as adding TTL outputs to the Raspberry Pi GPIO to trigger the start of an external optogenetics or photometry program, allowing for precise time locked videos. In addition, a logical next step for streamlining PiRATeMC is to convert the command line code into a unified Python code and/or a local graphical user interface (GUI) for an even more user friendly protocol.

Being able to easily record video of a large number of behavioral sessions simultaneously may usher in a new era of behavioral neuroscience analysis. In the field of drug addiction, we have been studying self-administration for decades using lever-pressing or nose-poking as the only proxy of ‘addictive-like behaviors’. By recording full behavioral sessions and analyzing the data with DeepLabCut and VAME, we have already begun to discover behavioral phenotypes that are highly predictive of an ‘addiction score’ that are not able to be detected by these standard behavioral metrics alone. Furthermore, behavioral neuroscience has long relied on human scoring and classification of behaviors, which suffers from serious confounding issues with inter-rater variability, the need for *a priori* knowledge of relevant behaviors to score, and capture of video data of high enough quality for accurate scoring. By using PiRATeMC in combination with machine-learning behavioral analysis, we address all three of these issues, and provide a method for unbiased, high-throughput behavioral analysis and classification.

We acknowledge that the use of this pipeline is still in its infancy and confounds still exist. First, PiRATeMC has only been validated using Linux and Mac operating systems and has not been tested using Windows. Another caveat is the difficulty in viewing the videos in real time while recording as alluded to above, although if this is required for behaviors without a predetermined timeline, either the 2.8” PiTFT touchscreen or 7” display (see **Supplemental Table 2**) can be used to view during recording with additional monetary cost, or the WebRTC platform can be used to view and record videos simultaneously at the cost of flexibility of recording parameters and synchronization of recording parameters. We hope the open-source nature of this pipeline will encourage users to expand this pipeline into many more interfaces beyond those outlined here, and further refine PiRATeMC to be optimized for each lab’s needs. Future editions of PiRATeMC will incorporate microcontrollers for closed-loop experiments (for example with optogenetic or MedAssociates equipment), and a streamlined data analysis pipeline whereby new video data is automatically detected and analyzed with a pre-set DeepLabCut model as soon as it finishes recording.

## Notes

https://github.com/alexcwsmith/PiRATeMC

### Competing Interest Statement

The authors have declared no competing interest.

### Summary of Updates

The primary purpose of this update is to include additional information on interfacing with third-party equipment. There are also several small clarifications within the text, and updated contact information (email) for the corresponding author.

