## Supplemental Tables 1 & 2 for "PiRATeMC: A highly flexible, scalable, and affordable system for obtaining high quality video recordings for behavioral neuroscience"

**Supplemental Information**

### Supplemental Table 1. Full list of nonessential accessories for PiRATeMC.

| Basic Camera Neccessities: | | | | |
|---|---|---|---|---|
| **Part:** | **Vendor (Suggested):** | **Product#:** | **Price** | **Notes:** |
| Raspberry Pi 3 Model B+ (or newer) | Adafruit Industries, Newark Electronics | 3775 (Adafruit), 49AC7637 (Newark), | $35.00 | Pi 3 B+ is convenient because it has standard HDMI ports and mini-USB power supply. Starting with Pi 4 there are only micro-HDMI ports and USB-C charging. The Pi 4 may be better if you want to apply computation or interface with RTCs (opto etc) |
| Raspberry Pi NoIR Camera Board v2 - 8MP | Adafruit Industries | 3100 | $29.95 | IR-sensitive camera - note the nomenclature here is super non-intuitive, the NoIR camera is the version that IS IR-sensitive, it is called NoIR because it does NOT have an IR filter. |
| Raspberry Pi Camera Board v2 - 8MP | Adafruti Industries | 3099 | | Camera version with an IR filter for recording under white light- same price as NoIR but didn't include price here so it's not included in total cost, as only one of the two is used. |
| 5V 2.5A Switching Power Supply with 20AWG MicroUSB Cable | Adafruit Industries | 1995 | $7.50 | Any mini-USB charging cable should work, but it needs to deliver a very stable 2.1A power, which many cheap low-quality cords do not, so I recommend buying this one. |
| Kingston 32GB microSD card | CDW | 5849358 | $4.99 | microSD card that serves as the main storage drive for the Raspberry Pi. |
| Flex Cable for Raspberry Pi Camera, or Display - 24" / 610mm | Adafruit Industries | 1730 | $2.50 | 24" cable is suitable for my applications, but you may want longer or shorter… options range from 2" to 2 meters. |
| | | **Total:** | **$79.94** | |

| Optional Camera Parts/Accessories: | | | | |
|---|---|---|---|---|
| **Part:** | **Vendor (Suggested):** | **Product#:** | **Price** | **Notes:** |
| 128GB SD Card for Expanded Storage Capacity | CDW | 6240103 | $29.99 | 128GB is probably far more than you'll ever need, as videos should only be temporarily stored here and are automatically transferred to a network location. |

| | | | | |
|---|---|---|---|---|
| Infrared light - JC Infrared Illuminator 4 Led High Power LED IR Array Illuminator | Amazon (or Fisher Sci) | https://www.amazon.com/Infrared-Illuminator-Power-Vision-Camera/dp/B01D73XM24 | $14.99 | Infrared light. Note power cord sold separately (though it is a common DC adapter). |
| Power Adapter for Infrared Light | Amazon (or Fisher Sci) | https://www.amazon.com/DC-Power-Adaptor-12-2-1MM/dp/B002O9QM5C/ | $5.64 | DC Adapter for IR light about. |
| PiTFT Plus Assembled 320x240 2.8", TFT + Resistive Touchscreen | Adafruit Industries | 2298 | $34.95 | One touch screen is convenient for setup, and if there is ever an issue with a Pi it is nice to be able to bring a display to it rather than having to plug into a monitor. |
| USB Battery Pack for mobility | Adafruit Industries | 1565 | $24.95 | Portable battery for experiments on-the-go. Lasts 6-12 hours depending on power consumption. |
| Case for Raspberry Pi | Adafruit Industries | 2604 | $8.96 | Not compatible with PoE hat. |
| Adafruit Raspberry Pi Camera Board Case with 1/4" Tripod Case with 1/4" Tripod | Adafruit Industries | 3253 | $2.95 | Camera Case |
| Adjustable Pi Camera Mount | Adafruit Industries | 1434 | $36.86 | Camera Mount |
| 7" Display | Adafruit Industries | 2718 | $79.95 | Allows watching video live while recording without connecting to monitor or reconfiguring PiTFT touchscreen. |

| Networking Equipment (for scaling up): | | | | |
|---|---|---|---|---|
| **Part:** | **Vendor (Suggested):** | **Product#:** | **Price** | **Notes:** |
| AWOW MiniPC | Amazon | https://www.amazon.com/dp/B07XBKFZVL | $209.99 | This is a MiniPC with two network ports that is very well suited for serving as the remote controller. Any computer can be used, but depending on your institutions IT policies, 2 network ports may be required. |
| NETGEAR 24-Port Gigabit Ethernet Unmanaged Switch, Plug-and-Play | CDW | JGS524NA | $119.99 | 24 Ports, each can run up to 23 cameras, with one port reserved for remote host controller. |
| Ethernet Cables | CDW | 2437674 | $12.99 | For 25' cable. You will want a range of lengths of cables, assuming some RPis will be further from your network switch than others. |
| Raspberry Pi PoE Hat | Adafruit Industries | 3953 | $20.00 | Allows Power over Ethernet, which is very convenient and reduces number of wires if scaling up to run man cameras via LAN. |
| 2-Port PCIe Network Card (not needed if you use AWOW MiniPC Above) | CDW | 3722710 | $124.99 | Most computers have only one ethernet port. If your institution IT department will not allow network switches (particularly cheap, unmanaged switches) on the backbone institutional network, or will not assign static IP addresses for all Pis, we provide instructions for configuring a remote host ('remote controller') to act as a DHCP server for the RPis, so that the remote controller can be connected to the backbone network (with internet connectivity), and will manage a separate LAN for the RPis. |
| 4-Port PCIe Network Card (not needed if you use AWOW MiniPC or 2-Port network card above) | CDW | 3877711 | $148.99 | See above, this expands ethernet capacity to 4 ports rather than 2. |

| Optional Microcontroller Interfacing Equipment: | | | | |
|---|---|---|---|---|
| **Part:** | **Vendor (Suggested):** | **Product#:** | **Price** | **Notes:** |
| GPIO Reference Card for ModelB+ Pi 2/3 | Adafruit Industries | 2263 | $2.50 | Makes identifying GPIO pins on the RPi easy to prevent having to look up the pin map every time you want to wire up a new configuration. |
| GPIO Expander and LED Driver | Adafruit Industries | 4886 | $4.95 | Suitable for an building a custom LED driver for a small fraction of the price of similar equipment marketed for optogenetics. |
| I2S MEMS Ultrasonic Microphone | Adafruit Industries | 3421 | $6.95 | Ultrasonic microphone capable of recoding up to 60kHz, enabling recoding of ultrasonic vocalizations from rodents. |
| General Purpose USB to GPIO/I2C/SPI | Adafruit Industries | 3421 | $6.95 | Allows control of GPIO, I2C, and SPI interfacing via USB output from RPi |
| Female/Male Jumper Wires (12"; Pack of 40) | Adafruit Industries | 824 | $7.95 | Jumper wires for microcontrollers |
| Female/Female Jumper Wires (12"; Pack of 40) | Adafruit Industries | 793 | $7.95 | Jumper wires for microcontrollers |
| Male/Female Jumper Wires (12"; Pack of 40) | Adafruit Industries | 793 | $7.95 | Jumper wires for microcontrollers |
| Creative Robotics & Interactive Construction Kit (CRIKIT) | Adafruit Industries | 3957 | $34.95 | Jumper wires for microcontrollers |

| | | | | |
|---|---|---|---|---|
| Peristaltic Pump | Adafruit Industries | 3910 | $24.95 | Peristaltic Pump with PWM capability for control of flow rate (could be used to build very cheap perfusion pump). |

**Supplemental Table 2. Full list of Raspivid options that can be added to recordVideo.sh line 3 (after raspivid)**

| Short form | Long form | Explanation |
|---|---|---|
| -? | --help | Help |
| -w | --width | Width of image (default 1920) |
| -h | --height | Height of image (default 1080) |
| -b | --bitrate | Bits per second |
| -o | --output | Output filename |
| -v | --verbose | Output verbose info during run |
| -t | --timeout | Time before takes picture and shuts down |
| -d | --demo | Semo mode- cycle through camera options, no capture |
| -fps | --framerate | Number of frames to record per second |
| -e | --penc | Display preview image after encoding |
| -p | --preview | Preview window settings |
| -f | --fullscreen | Fullscreen preview mood |
| -n | --nopreview | Do not display preview window |
| -sh | --sharpness | Image sharpness (-100 to 100) |
| -co | --contrast | Image contrast (-100 to 100) |
| -br | --brightness | Image brightness (0 to 100) |

| | | |
|---|---|---|
| -sa | --saturation | Image saturation (-100 to 100) |
| -ISO | --ISO | Set capture ISO- brightens/darkens based on light level |
| -vs | --vstab | Turn on video stabilization |
| -ev | --ev | Exposure compensation (adjusts lighter or darker) |
| -ex | --exposure | Set exposure mode (options= off, auto, night, nightpreview, backlight, spotlight, sports, snow, beach, verylong, fixedfps, antishake, fireworks |
| -awb | --awb | Set auto white balance (options= off, auto, sun, cloud, shade, tungsten, fluorescent, incandescent, flash, horizon, greyworld) |
| -ifx | --imfx | Set image effects (options= none, negative, solarise, sketch, denoise, emboss, oilpaint, hatch, gpen, pastel, watercolour, film, blur, saturation, colourswap, washedout, posterise, colourpoint, colourbalance, cartoon) |
| -cfx | --colfx | Set color effect (U:V) |
| -mm | --metering | Set metering mode (average, spot, backlit, matrix) |
| -rot | --rotation | Set image rotation (0-359) |
| -hf | --hflip | Set horizontal flip |
| -vf | --vflip | Set vertical flip |

**Supplemental links and resources:**
[https://github.com/alexcwsmith/PiRATeMC](https://github.com/alexcwsmith/PiRATeMC)
https://thepihut.com/blogs/raspberry-pi-roundup/raspberry-pi-camera-board-raspivid-command-list
